# Towards the Rational Design of RsmE Small-RNA Binders: Insights from Molecular Dynamics Simulations

**DOI:** 10.1101/2025.10.23.684185

**Authors:** Agustín Ormazábal, Juliana Palma, Gustavo Pierdominici-Sottile

## Abstract

The RsmZ–RsmE interaction is a key element in post-transcriptional regulation in *Pseudomonas* species. Experimental studies have shown that alternative fragments of RsmZ, despite displaying only subtle sequence and structural differences, exhibit markedly distinct binding affinities for RsmE. To complicate matters further, the affinities measured for isolated fragments differ substantially from those observed when the same segments are embedded in the full-length sRNA molecule. To explore the origin of these discrepancies, we generated computational models of RsmE dimers bound to one or two sRNA stem loops, including several experimentally studied variants, a couple of truncated forms, and a synthetic construct created by linking two native stem loops with an also native single-stranded region. The unbinding of the RNA fragments from these complexes was studied using Umbrella Sampling simulations, which revealed that base pairs located in the stems, as well as the presence of a linker region, shape the interaction landscape between the protein and RNA, thereby modulating binding affinities. Thus, our findings offer a structural and mechanistic basis for interpreting the experimentally observed differences among RsmZ fragments and establish a conceptual foundation for the future computational design of synthetic sRNAs capable of fine-tuning the RsmZ/RsmE regulatory system in a predictable manner.

## 1 Introduction

Within the vast diversity of microorganisms, plant-associated species rely on the synthesis of secondary metabolites to adapt and thrive in their immediate environment.^1^ In particular, certain rhizosphere-dwelling bacteria, such as Gram-negative fluorescent *Pseudomonas* spp., have been identified as biocontrol agents capable of protecting crop plants against phyto pathogenic fungi, bacteria, and nematodes through the production of secondary metabolites.^2,3^ In *Pseudomonas protegens*, one such metabolite is hydrogen cyanide (HCN), whose biosynthesis is regulated in a cell density-dependent manner. ^4,5^ HCN plays a key ecological role due to its proposed contribution to biotic interactions. ^6^ Accordingly, the use of HCN-producing bacteria as biopesticides appears as an eco-friendly strategy for sustainable agriculture. The biosynthesis of HCN is directed by the *hcnABC* operon, which encodes the HCN synthase enzyme complex. Specifically, the *hcnA* mRNA encodes the A subunit of this trimeric complex. ^7^

For decades, transcription initiation was considered the primary regulatory stage of gene expression. More recently, however, the translational regulation of mRNAs in prokaryotes has attracted increasing attention.^8^ In this context, non-coding small RNAs (sRNAs) have emerged as key players in modulating gene activity,^9^ further highlighting the importance of translational control in prokaryotes. A paradigmatic example is the Csr/Rsm system, a broadly conserved post-transcriptional regulatory mechanism in *Eubacteria*, which enables cells to fine-tune metabolism in response to environmental signals. ^8,10–13^ Its importance is underscored by its occurrence in roughly 75% of bacterial species, ^14^ with more than two thousand species annotated in the Pfam database (PF02599) as harboring this system.^15^ In *E. coli*, nearly 20% of all mRNAs are estimated to be regulated through this pathway. ^7,16–18^

One of the central components of the Csr/Rsm system is the protein CsrA (from its acronym <u>C</u>arbon <u>S</u>torage <u>R</u>egulator “A”). In *Pseudomonas protegens*, the CsrA homologue is known as RsmE. ^7,19^ Hereafter, we will use this name to refer to the protein, as it is the one employed in our computational models. RsmE binds to A(X)GGAX motifs located in the 5’ <u>u</u>ntranslated <u>r</u>egion (5’ UTR) of target mRNAs, preventing access of the 16S ribosomal subunit and thereby repressing translation. In the motif, X denotes any nucleotide, while the parentheses indicate an optional position. ^20–24^ Structurally, RsmE is a homodimer whose core is formed by intertwined *β*-strands from both subunits. Two identical RNA-binding sites are positioned on opposite sides of the protein. ^20–24^ Depending on the structural and sequence context of the A(X)GGAX motifs within the RNA, their affinity for RsmE can vary by more than five orders of magnitude. ^25^

In *Escherichia coli* and *Pseudomonas* spp., several mRNAs regulated by CsrA/RsmE contain multiple putative binding motifs upstream of their Shine–Dalgarno sequences. For example, *hcnA* carries four A(X)GGAX motifs in this region. ^3^ A major breakthrough in understanding the regulatory mechanism of RsmE was the elucidation of the three-dimensional structure of its complex with a 20-nucleotide RNA construct derived from the *hcnA*5 ^*′*^-UTR. This construct contains the A(X)GGAX motif closest to the translation initiation codon. ^26^ The structure revealed that the motif (ACGGAU in this case) is located at the apex of a stem loop (SL) stabilized by a three–base-pair stem formed by adjacent nucleotides (see Figure 1). It also showed that RsmE makes specific contacts with all nucleotides of the hexanucleotide loop, in addition to nonspecific interactions with the C7–G14 and U6–A15 base pairs. ^26^

**Figure 1.**
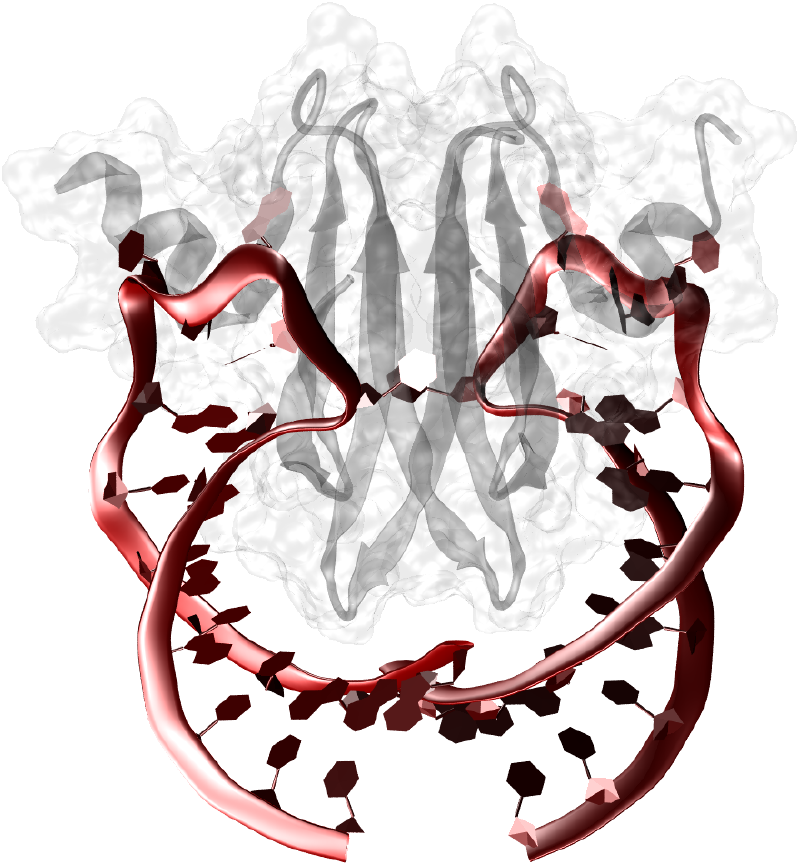
Pictorial representation of the complex formed by an RsmE dimer bound to two SLs corresponding to the 20-nt segment derived from the *hcnA* mRNA. This structural arrangement is conserved in complexes formed between RsmE and other RNA stem loops.

In parallel to the protein CsrA/RsmE and its cognate mRNAs, this regulatory system relies on a specific class of sRNA as essential counterparts. These sRNAs also contain A(X)GGAX motifs that sequester CsrA/RsmE proteins from their target mRNAs through competitive binding, thereby relieving translational repression.^27–36^ The intrinsic flexibility of the RNAs involved, together with their multiple binding motifs and the two RNA-binding sites in the CsrA/RsmE dimers, gives rise to a remarkably intricate regulatory mechanism at the molecular level. Owing to this complexity, and to its pivotal role in controlling the expression of diverse metabolic pathways across a wide range of bacterial species, ^10,24,37–40^ numerous investigations have sought to uncover the principles underlying its functioning.^20,24–26^

Among the experimental investigations, structural studies of the complexes formed by Csr/Rsm proteins and RsmZ fragments have been particularly illuminating. ^24,25,41^ RsmZ is one of the non-coding sRNA that participates in the Csr/Rsm regulatory system in *Pseudomonas*. It contains eight A(X)GGAX binding motifs. Five of them are located at the apices of stem loops SL1 to SL5, resembling the one described for the 20-nt construct of *hcnA*. Another motif, relevant for the current discussion, lies in an 8-nt single-stranded region (SSR) connecting SL2 and SL3. Duss and co-workers characterized the complexes generated by RsmE and alternative fragments of the *Pseudomonas fluorescens* RsmZ. In 2014, they reported the nuclear magnetic resonance structures of stem loops SL1 to SL4 bound to RsmE, as well as the structure of the complex formed by the binding motif located in the 8-nt SSR. ^25^ All these complexes exhibit twofold symmetry, with one RNA segment bound on each side of the RsmE dimer, similar to that shown in Fig. 1 for the 20-nt construct from *hcnA*.

Subsequently, the same group reported the structure of the complex formed by three RsmE dimers and a 72-nt RsmZ fragment spanning SL1 to SL4.^24^ The study also analyzed NMR chemical shift perturbations observed during titration of RsmZ(1-89) with increasing stoichiometric amounts of ^2^H-^15^N-labeled RsmE, aimed to determine the order in which the different binding motifs are occupied. The results revealed that complex formation proceeds through a sequential mechanism: the first dimer binds simultaneously to SL2 and SL3, the second dimer binds to SL1 and SL4, and the third binds to the AGGAX motif located in the SSR between SL2 and SL3, as well as to a site located beyond nucleotide 72. More recently, Jia and co-workers applied cryo-EM to solve the structure of the full-length RsmZ from *Pseudomonas aeruginosa* in complex with three RsmA units (a homolog of RsmE).^41^ Interestingly, in this case, sequential binding was not observed, and the pair of binding motifs engaged with each protein dimer differed from those reported in Ref. 24. Specifically, one dimer was embraced by SL1 and SL5, another by SL2 and SL3, and the third by SL4 and the AGGAX motif sited in the SSR that links SL2 with SL3.

Also crucial for advancing our understanding of the Csr/Rsm regulatory system was the determination of the dissociation constants, *K*_*d*_, for the different complexes mentioned in the previous paragraph. In this regard, the first unexpected finding came from measuring *K*_*d*_ for the individual stem loops of RsmZ. ^25^ The study revealed that only SL2 displays an affinity for RsmE comparable to that of the 20-nucleotide fragment of *hcnA*. The other stem loops, as well as the binding motif in the SSR, showed *K*_*d*_ values one or two orders of magnitude higher. With such low constants, it is not clear how RsmZ could compete with the mRNA for the protein. Another striking observation was that, whereas the low-affinity stem loops are characterized by a single dissociation constant, the complexes involving SL2 exhibit two distinct constants, similar to what had been reported for the 20-nt *hcnA* fragment. Thus, the reported *K*_*d*_ for the dissociation of SL2 from RsmE-(SL2)_2_ is 185 nM, whereas its dissociation from RsmE-(SL2)_1_ occurs with the much lower *K*_*d*_ of 16 nM. Although plausible hypotheses were proposed to explain the correlation between strong binding and the presence of two dissociation constants, no direct evidence has yet confirmed them.

The apparent inconsistencies between the relatively weak affinities of most RsmZ stem loops and the stronger affinity of *hcnA* were resolved once the dissociation constants were measured for the 72-nt RsmZ fragment.^24^ Unlike the experiments described above, in which two identical RsmZ fragments bind symmetrically to both sides of an RsmE dimer, full-length RsmZ engages each RsmE dimer through binding motifs located in different stem loops or in the SSR. These experiments revealed *K*_*d*_ values of 150 nM for RsmE bound to SL2/SL3, 100 nM for SL1/SL4, and 200 nM for the SSR motif together with the motif located downstream of nucleotide 72. Thus, the experiment demonstrated that the affinities of the individual motifs are determined not only by their intrinsic properties but also, to a significant extent, by their molecular context. These findings clarified how RsmZ effectively competes with *hcnA* for RsmE binding. Nevertheless, the mechanistic basis for the observed increase in *K*_*d*_ remains unresolved. Recent experiments conducted on full-length RsmZ from *Pseudomonas aeruginosa* ^41^ reported dissociation constants of the same order of magnitude as those for RsmZ from *Pseudomonas fluorescens*, albeit slightly higher.

Molecular dynamics simulations have provided a plausible explanation for the sequential binding of RsmE to the 72-nt RsmZ fragment.^15^ They revealed that the alternative binding motifs follow a well-defined pattern. When the fragment is unbound, only the motifs that recruit the first RsmE dimer are accessible, while the other three remain occluded. Once the first dimer is bound, the motifs that recruit the second dimer become exposed, whereas the site for the third dimer remains permanently inaccessible. Finally, after two RsmE dimers are bound, the last motif becomes available. The study also found that the two motifs that engage a single RsmE dimer are not exposed at the same time. This feature may play a role in preventing two distinct dimers from simultaneously occupying binding motifs intended for a single dimer. From this, it follows that RsmE initially interacts with a single motif, after which the RNA fragment undergoes a conformational rearrangement that enables it to engage the dimer from the other side.

More recently, MD simulations were employed to reveal the details of how an RsmE dimer binds to the motif located in the SSR of RsmZ that links SL2 with SL3. ^42^ The study not only described how the interactions between the molecular fragments are established from the initial stages of their mutual recognition, but also highlighted the crucial role of the SSR’s flexibility, which allows it to deform and approximate the shape of a stem loop when pulled by the RsmE dimer. In this article, we present the results of a molecular dynamics simulation study investigating the binding mechanism between alternative RsmZ fragments and an RsmE dimer. These simulations were designed to elucidate the molecular determinants underlying the variable behavior observed for these fragments in the experimental studies discussed above. By integrating the insights from our analyses, we rationalize how seemingly small sequence or structural differences modulate binding behavior. Together, these findings lay the groundwork for the computational design of sRNA molecules capable of modulating the Csr/Rsm regulatory mechanism in a predictable manner.

## 2 Methods

### 2.1 Model Structure Construction

CsrA/RsmE is a symmetrical homodimer stabilized by intertwined *β*-sheets. Each monomer contains five *β*-strands ( *β*1– *β*5) and a short *α*-helix. There are two symmetrical RNA-binding surfaces located on opposite sides of the dimer, each capable of recognizing an A(X)GGAX RNA motif. In this work, eleven computational models were generated based on previously reported structures of RsmE bound to different RNA constructs. As starting templates, we used Protein Data Bank entries 2MFE, 2MFF, and 2JPP. All these structures feature two identical RNA stem loop (SL) fragments bound to an RsmE dimer. Specifically, 2MFE and 2MFF correspond to RsmE in complex with two copies of SL2 and SL3 from RsmZ, respectively, ^25^ whereas 2JPP contains two copies of the 20-mer construct derived from the *hcnA* transcript.^26^ We note, however, that the RNA fragments in these PDB structures do not exactly match their sequences in RsmZ. In the experimental constructs, two or three additional complementary nucleotides were introduced to stabilize stem formation in the corresponding stem loops. Therefore, from this point onward, we refer to these stem loops as the “experimental SLs”.

Three computational models were directly derived from the above-mentioned PDB structures. These are referred to as RsmE-(SLX)_2_, where X = 2 or 3 denotes the corresponding RsmZ stem loop, while we will use X = 0 to denote the stem loop derived from the *hcnA* transcript, for the sake of conciseness. Another set of three models, referred to as RsmE-(SLX)_1_, was generated from the RsmE-(SLX)_2_ models by removing one of the SLs. Note that, as the PDB structures are symmetrical, it is immaterial which stem loop was deleted.

Five additional *in-silico* tailored models were generated to investigate the contribution of stem-loop nucleotides to binding strength, as well as the impact of the linker region connecting RNA fragments. Two of these models, hereafter referred to as RsmE-(SL2^(s)^)_1_ and RsmE-(SL2^(s)^)_2_, contain a shortened version of the experimental SL2 in which nucleotides G1–G3 and U18–C20 were removed. Another pair, named as RsmE-(BM)_1_ and RsmE-(BM)_2_, correspond to complexes with one or two copies, respectively, of the 6-nt binding motif present in the 20-nt *hcnA* construct. In other words, those complexes were built by deleting the whole stem (nucleotides G1–C7 and G14–C20) from the mRNA fragment evaluated in the experiments of Schubert et. al. ^26^ Finally, the last model, designated RsmE-(SL2^(l)^)_2_, comprises two full-length SL2 elements connected through the single-stranded SL2–SL3 linker region of RsmZ (nucleotides 36–43). To construct this model, SL2 and SL3 from the 72-nt RsmZ structure^24^ were aligned with the two SL2 fragments of RsmE-(SL2)_2_, and the coordinates of the linker region were extracted and incorporated into a new PDB file with a modified topology to ensure sequence continuity.

The connections were designed to preserve the 5’–3’ orientation of both SLs in the final chain. A schematic representation of all models analyzed in this work is shown in Figure 2.

**Figure 2.**
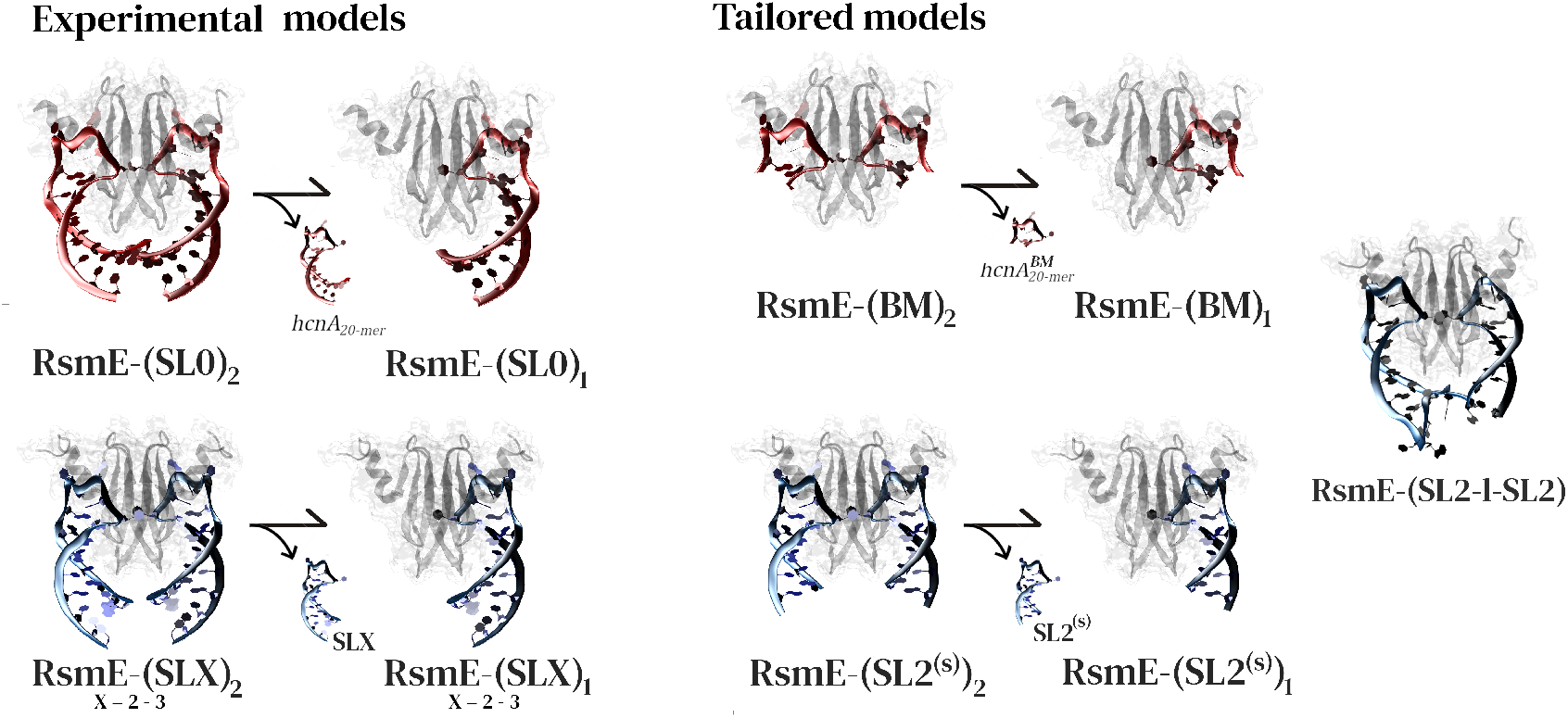
Graphic summary of the models studied in this work, with a schematic illustration of how they were constructed.

### 2.2 Systems Preparation and General MD Settings

We employed the following protocol to generate the computational models used in the MD simulations. The initial structures in PDB format were inserted into the LEaP module of AMBER18.^43^ Each system was solvated in an octahedral box of water molecules, with the box walls placed 36 Å away from the closest solute atom. This large box size was required because the RNA fragments adopt extended conformations when pulled out of the complex during the Umbrella Sampling (US) simulations. Protein and ion interactions were described with the ff14SB force field,^44^ RNA molecules with frcmod.ROC-RNA,^45^ and water with the TIP3P model. ^46^ All simulations were carried out under periodic boundary conditions. The Particle Mesh Ewald method^47,48^ with a real-space cutoff of 9.0 Å was employed to compute the electrostatic interactions. The cutoff for the rest of the non-bonded interactions was also set to 9.0 Å. We implemented the SHAKE algorithm to constrain the lengths of bonds involving hydrogen atoms, thus allowing using a time step of 2.0 fs in the simulations. All simulations were performed with the PMEMD module of AMBER18.

The models generated with the LEAP package were first energy-minimized and subsequently heated under NVT conditions for 2 ns to reach 303 K. Temperature control was achieved using a Langevin thermostat with a collision frequency of 1.0 ps^−1^.^45^ The heating phase was followed by a 20 ns equilibration under NPT conditions, allowing the system density to relax. Pressure was regulated with a Berendsen barostat using a coupling constant of 2.0 ps. The final structures of the equilibration stages were used as starting configurations for the US calculations.

### 2.3 Umbrella Sampling Simulations

The Umbrella Sampling technique was employed to simulate the unbinding of a single RNA fragment from complexes with one or two copies of it. Recall that, in US simulations, all coordinates except the reaction coordinate are assumed to be in equilibrium. In this framework, reversing the SLX unbinding from RsmE-(SLX)_2_ represents SLX binding to a half-occupied dimer, while reversing SLX unbinding from RsmE-(SLX)_1_ represents its binding to an empty dimer. Similarly, we applied US to dissociate each SL2 from the 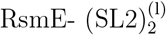 complex in a stepwise manner.

In all cases, the reaction coordinate 𝒳 was defined as the distance between two centers of mass (COMs). The COM of the RNA construct was calculated using the heteroatoms of the A(X)GGAX motif together with the flanking nucleotides forming the first 3–4 base pairs of the hairpin surrounding the binding motif. The protein COM was computed from the C_*α*_ atoms of residues 1–7 (corresponding to *β*1_*A*_) and residues 89–104 (forming *β*4_*B*_, *β*5_*B*_, and the connecting loop). This protein region defines the dimer binding surface. A schematic representation of the regions used to define 𝒳 is shown in Fig. 3.

**Figure 3.**
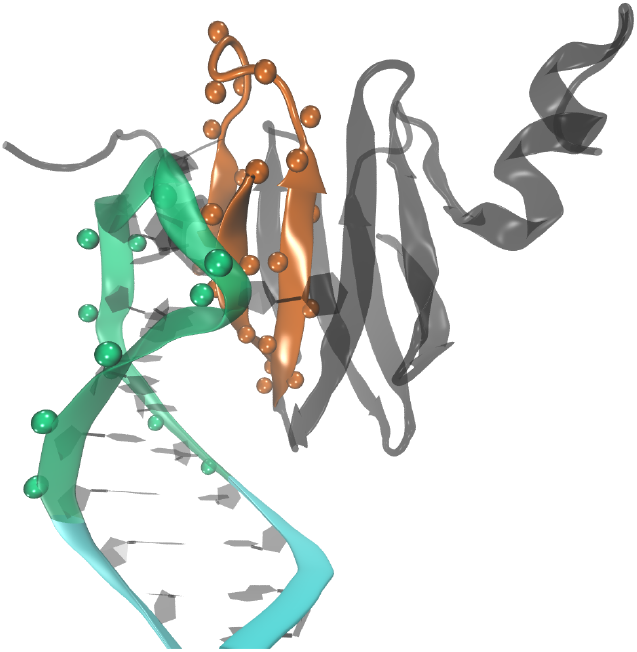
Structural regions used to define the reaction coordinate 𝒳. The coordinate corresponds to the distance between the centers of mass (COMs) of two groups: (i) the heteroatoms of the A(X)GGAX motif together with the flanking nucleotides forming the first 3–4 base pairs of the hairpin surrounding the binding motif (green spheres), and (ii) the C_*α*_ atoms of strands *β*1_*A*_, *β*4_*B*_, and *β*5_*B*_, together with the loop connecting *β*4_*B*_ and *β*5_*B*_ (orange spheres). This protein region constitutes its binding surface.

To capture the entire unbinding event, the reaction coordinate 𝒳 was increased from its value in the initial structure (the last frame from the equilibration stage) to approximately 25–30 Å. The spacing between adjacent window centers was set to 0.1 Å, and the final structure from window *i* served as the starting point for simulations in window *i*+1. Each window was sampled for 10 ns, with 𝒳 recorded every 0.1 ns. The force constant of the harmonic bias potential was set to 350.0 kcal/molÅ^2^. To enable structural analysis throughout the unbinding process, snapshots were saved every 0.2 ns within each window. In total, the simulations amounted to 1.3–1.8 *µ*s per model.

### 2.4 Analysis

All the analyses described in this section were performed with module CPPTRAJ of AMBER24. Equilibration and stability of the computational models used to start the US simulations were assessed by analyzing time evolution of the RMSD of the complex structures, as well as the number of contacts and H-bonds between the molecular partners. Also, the number of contacts and H-bonds were analyzed with the snapshots from each window of the Umbrella Sampling simulations.

The potential of mean force (PMF) for each (un)binding process was computed from the US data using three alternative protocols: the Weighted Histogram Analysis Method (WHAM), ^49^ the Dynamic Histogram Analysis Method (DHAM), ^50^ and the Bennett Acceptance Ratio (BAR).^51^ Custom FORTRAN codes were employed for these calculations. Since the three approaches rely on different assumptions, the agreement among their outcomes increases confidence in the resulting PMF profiles. For WHAM and DHAM, the reaction coordinate was binned into 0.025 Å segments. Convergence of the WHAM algorithm required 700000 iterations. DHAM could not be applied across the entire range of the reaction coordinate, but yielded reliable results for selected segments, which were then assembled to construct the complete curve. For BAR, 500 iterations were performed to estimate the free energy differences between adjacent distributions. Several tests were carried out to examine the convergence and internal consistency of the PMFs, as well as to estimate their statistical uncertainty. They are presented and discussed in the Supplementary Information section.

### 2.5 Entropic contributions: quasi-harmonic approximation (QHA)

Entropy changes play a significant role in explaining the different strengths with which SL2 binds to RsmE in the complexes where the SL2 fragments are either separated, RsmE-(SL2)_2_, or connected through a linker region, RsmE-(SL2^(l)^)_2_. To quantify this effect, we employed the quasi-harmonic approximation to estimate the entropy of the reactants and products in both processes, under the assumption that the dominant contribution arises from the configurational entropy of the molecules. The two processes considered for comparison are:

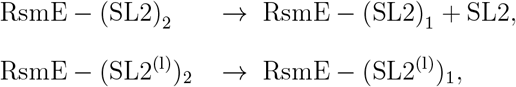

where RsmE-(SL2^(l)^)_1_ denotes the complex in which one of the stem loops of the (SL2^(l)^)_2_ construct is detached from its binding surface on RsmE, while the other remains bound. The entropy changes of these processes were estimated from the QH-entropies of the intervening molecules as,

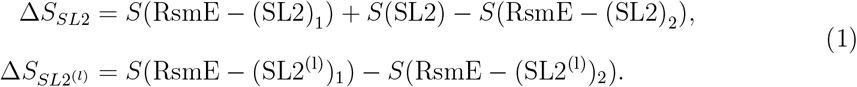

To carry out the QHA evaluations we performed 40 independent 50-ns MD simulations for RsmE-(SL2)_2_, RsmE-(SL2)_1_, SL2, RsmE-(SL2^(l)^)_2_, and RsmE-(SL2^(l)^)_1_. Then, all the simulations for each system were combined to form a concatenated trajectory used in the QHA computations. This procedure accelerates the convergence of the QH modes by reducing the effect of spurious correlations that alter the eigenvalues (and eigenvectors) of the covariance matrices. ^52,53^ Overall translation and rotation were removed from the concatenated trajectories, and the atomic displacements of all heavy atoms were computed with respect to the average structure. Finally, the mass-weighted covariance matrix of these displacements was constructed and diagonalized to determine the eigenvalues *{λ*_*i*_ *}* corresponding to the quasiharmonic modes. Under the QHA, each mode is treated as an effective harmonic oscillator whose classical entropy is evaluated as,

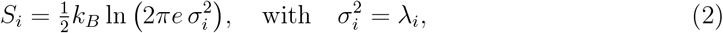

Accordingly, the total configurational entropy of each molecule was approximated as *S* ≈ Σ_*i*_ *S*_*i*_, where the summation extends to all degrees of freedom of the system. These entropies were introduced in Eq. 1 to determine the entropy change upon unbinding. The convergence of the entropy estimations with respect to the number of individual trajectories included in the concatenated one is presented in Fig. **??** of the Supplementary Information section.

## 3 Results

### 3.1 Simulations Mirror Experiments

Figure 4 presents the Potential of Mean Force (PMF) profiles characterizing the unbinding of SLX, from the RsmE-(SLX)_1_ and RsmE-(SLX)_2_ complexes, for the stem loop formed by the 20-nt *hcnA* fragment (SL0), as well as for SL2 and SL3 of RsmZ. From these profiles, the Free Energies of Binding (FEB) were estimated as the PMF plateau reached at sufficiently large reaction coordinate values. The resulting FEBs for SLX binding to RsmE (FEB_1_) and to RsmE-(SLX)_1_ (FEB_2_) are listed in Table 1, alongside the experimentally determined dissociation constants ( *K*_*d*_).

**Table 1.** Estimated free energy barriers (FEB) for each dissociation event in complexes RsmE-(SLX)_1_ and RsmE-(SLX)_2_, contrasted with their experimental dissociation constants *K*_*d*_. The last three columns indicate the number of nucleotides in the stem, loop, and the whole RNA construct, respectively. We recall SL0 is a compact notation we introduced to denote the SL formed by the 20-nt fragment of *hcnA*.

| | FEB<sub>1</sub><br>(Kcal/mol) | FEB<sub>2</sub><br>(Kcal/mol) | $K_{d1}$<br>(indiv) | $K_{d2}$<br>(indiv) | $K_{d1,2}$<br>(pair) | nt in stem<br>(5'+3') | nt in<br>loop | Total nt |
| --- | --- | --- | --- | --- | --- | --- | --- | --- |
| SL0 | 71.34±4.54 | 37.14±6.00 | 0.020 | 0.13 | NA | 2×7 | 6 | 20 |
| SL2 | 60.45±6.58 | 37.70±3.09 | 0.016 | 0.185 | 0.10 | 2×(4+4) | 6 | 22 |
| SL3 | 31.0±4.68 | 28.0±2.78 | 1.5 | 1.5 | 0.10 | 2×(4+4) | 5 | 21 |

**Figure 4.**
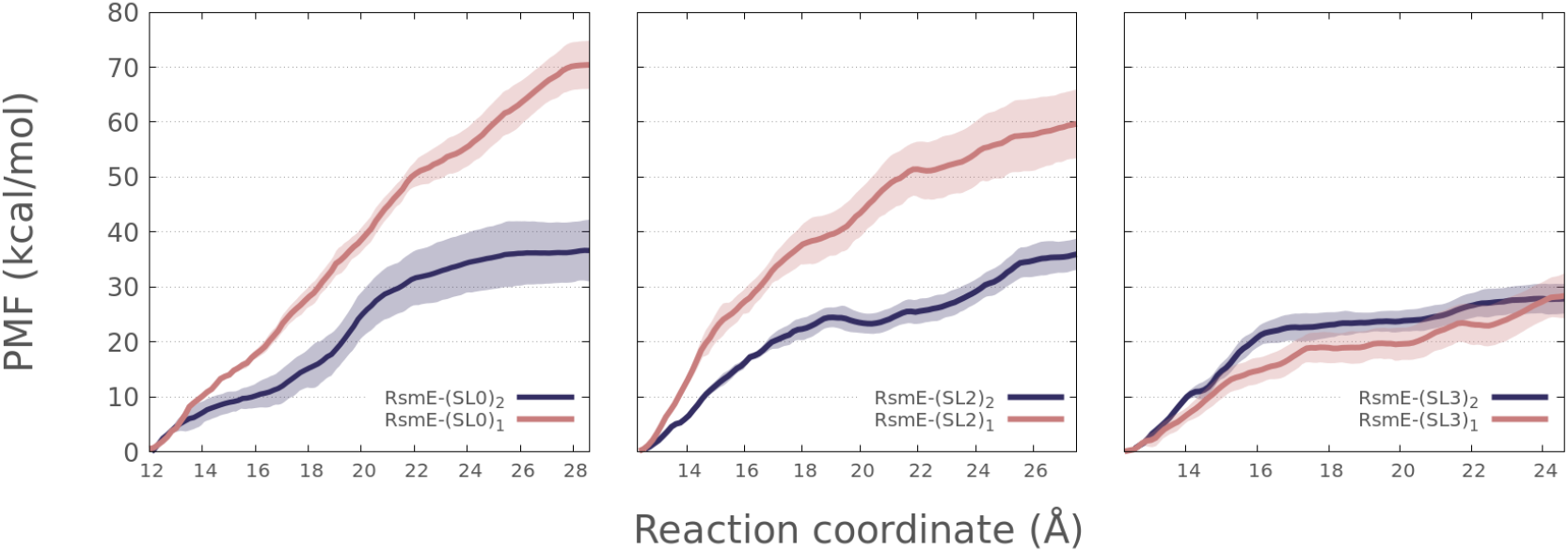
Potentials of Mean Force (PMFs) describing the detachment of native stem loops (SLX) from RsmE-(SLX)_1_ and RsmE-(SLX)_2_ complexes. The left panel reports the PMF for the stem loop formed by the 20-nt fragment derived from *hcnA* (denoted SL0 in the figure). The middle and right panels show results for the SL2 and SL3 fragments of RsmZ from *Pseudomonas fluorescens*, respectively. Shadows behind the curves indicate their estimated statistical uncertainty.

Analysis of Fig.4 and Table 1 reveals a clear agreement between experimental and computational trends. Thus, the calculated FEBs for SL3 dissociation from RsmE-(SL3)_2_ and RsmE-(SL3)_1_ are nearly indistinguishable. In contrast, SL2 and the *hcnA*-derived stem loop display markedly different behavior, with FEB_1_ consistently exceeding FEB_2_. These computational observations align well with the experimentally determined dissociation constants ( *K*_*d*_). While a single *K*_*d*_ has been reported for SL3 dissociation from both RsmE-(SL3)_2_ and RsmE-(SL3)_1_,^24,25^ two distinct *K*_*d*_ values were measured for SL2 and for SL0, being the *K*_*d*_ for RsmE-(SLX)_2_ systematically larger than that of RsmE-(SLX)_1_. This agreement between calculations and experiments suggests that the models employed in the simulations are able to capture the key features of the systems under study, lending confidence to the explanations and interpretations derived from them, which are presented below.

### 3.2 Variations in Protein-RNA Contacts Underlie FEB Differences

A first clue as to why two distinct FEBs are observed for the dissociation of SL2 and the SL derived from the *hcnA* construct comes from the analysis of movie S1. This movie was generated from snapshots of the US simulation for the SL2 detachment from RsmE-(SL2)_2_. By reversing the order of the frames, the movie effectively illustrates the binding process of SL2 to the RsmE-(SL2)_1_ complex. The visualization reveals that molecular recognition begins at the upper region of the RNA fragment, where the GGA motif engages with the RsmE binding surface. Once this initial contact is established, the stem progressively forms its own interactions, starting from the upper portion and propagating downward. Toward the end of the process, the lowest base pairs of the incoming SL2 attempt to accommodate themselves into position. However, the spatial region they require is occupied by the lowest base pairs of the SL2 already bound. As a result, the incoming RNA fragment must partially displace the pre-bound SL2, weakening the interactions of the latter with RsmE. The energetic cost of this displacement, together with the reduced capacity of the incoming SL2 to form interactions as strong as those observed when only a single fragment is bound, would explain the significantly weaker FEB measured for the second binder. An analogous situation is observed for the SL formed by the 20-nt *hcnA*fragment.

To provide more robust support for the pictorial explanation presented above, we quantified the number of contacts between RsmE and the RNA fragments in the RsmE-(SLX)_1_ and RsmE-(SLX)_2_ complexes. Because protein-RNA contacts are present in both cases and their numbers fluctuate across samples, we characterized the differences by computing probability distribution functions for the contact counts. For the results presented below, we considered a residue-nucleotide contact existed when the distance between not hydrogen atoms was shorter than 4.2 Å. Qualitatively similar results are obtained for a 3.5 Å and a 4.5 Å distance cut-off. The middel panel of Fig. 5 shows the results for the complexes formed by SL2. It is observed that the two distributions partially overlap, but the one corresponding to the RsmE-(SL2)_1_ complex is clearly shifted to the right of the one corresponding to RsmE(SL2)_2_. Thus, the most probable number of contacts is 312 for RsmE-(SL2)_1_ and 280 for RsmE-(SL2)_2_. As the distributions are unimodal and rather symmetric, these numbers also agree with the average number of contacts for each complex. The probability distribution functions for the complexes containing the 20-nt stem loop derived from *hcnA* present the same characteristics, but are slighted shifted to the right of those corresponding to SL2, as can be seen in the left panel of Fig. 5. This difference agrees with the larger FEBs for these *hcnA* complexes, as well as their smaller experimental *K*_*d*_ values (see Fig.4 and Table 1).

**Figure 5.**
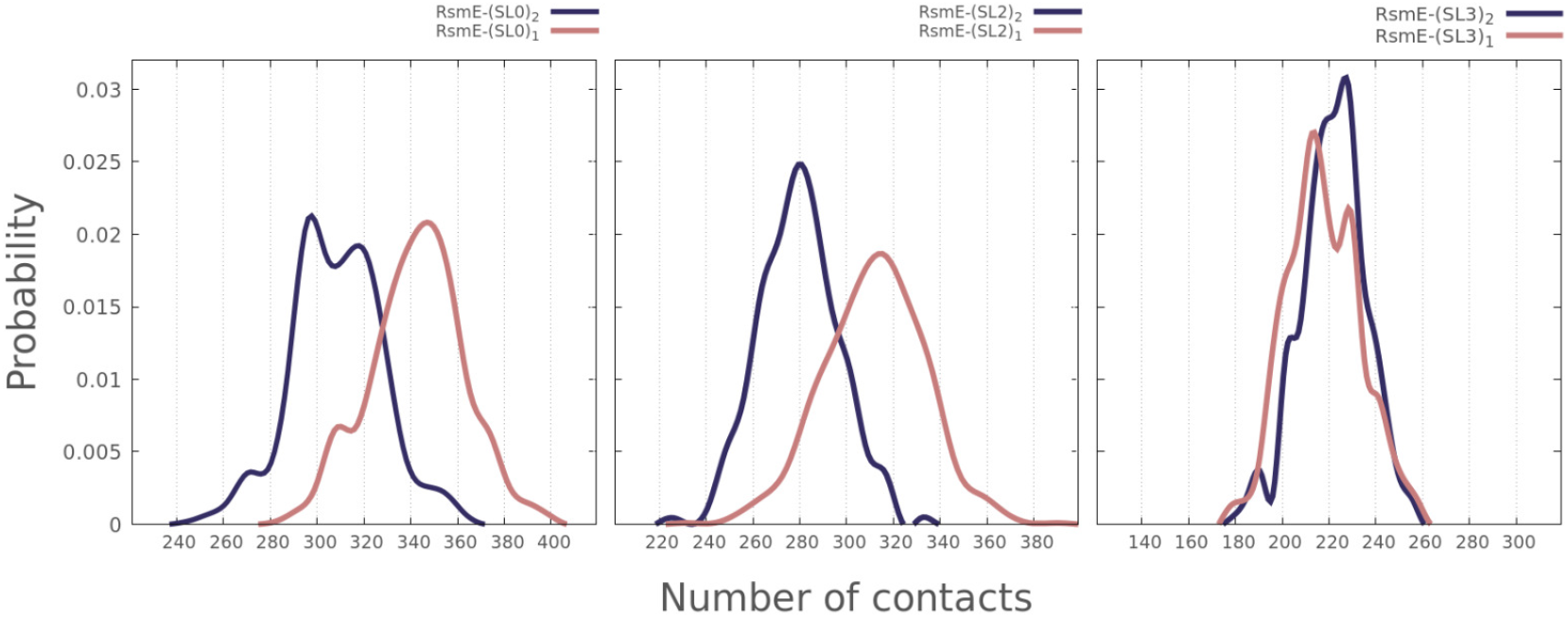
Probability distribution functions for the number of RNA–protein contacts in complexes formed by the 20-nt fragment of *hcnA* (SL0, left panel), SL2 of RsmZ (middle panel), and SL3 of RsmZ (right panel).

### 3.3 Stem-Base Repulsion Reduces Protein–RNA Contacts

Having shown that, in the cases of SL0 and SL2, the differences in FEBs originate from a higher number of protein–RNA contacts in single-SL complexes compared to double-SL ones, we next sought to identify the molecular determinants underlying this difference. To this end, we constructed the RsmE-(SL2^(s)^)_1_ and RsmE-(SL2^(s)^)_2_ models. As described in Section 2.1, they were generated by removing the three lowest base pairs from the SL2 experimentally characterized in Ref. 25. The PMFs for the detachment of SL2^(s)^ from both complexes are shown in the left panel of Fig. 6. It can be seen that the two profiles nearly overlap. Besides, they closely resemble the PMF for the detachment of SL2 from RsmE-(SL2)_1_, also shown in the left panel of the figure.

**Figure 6.**
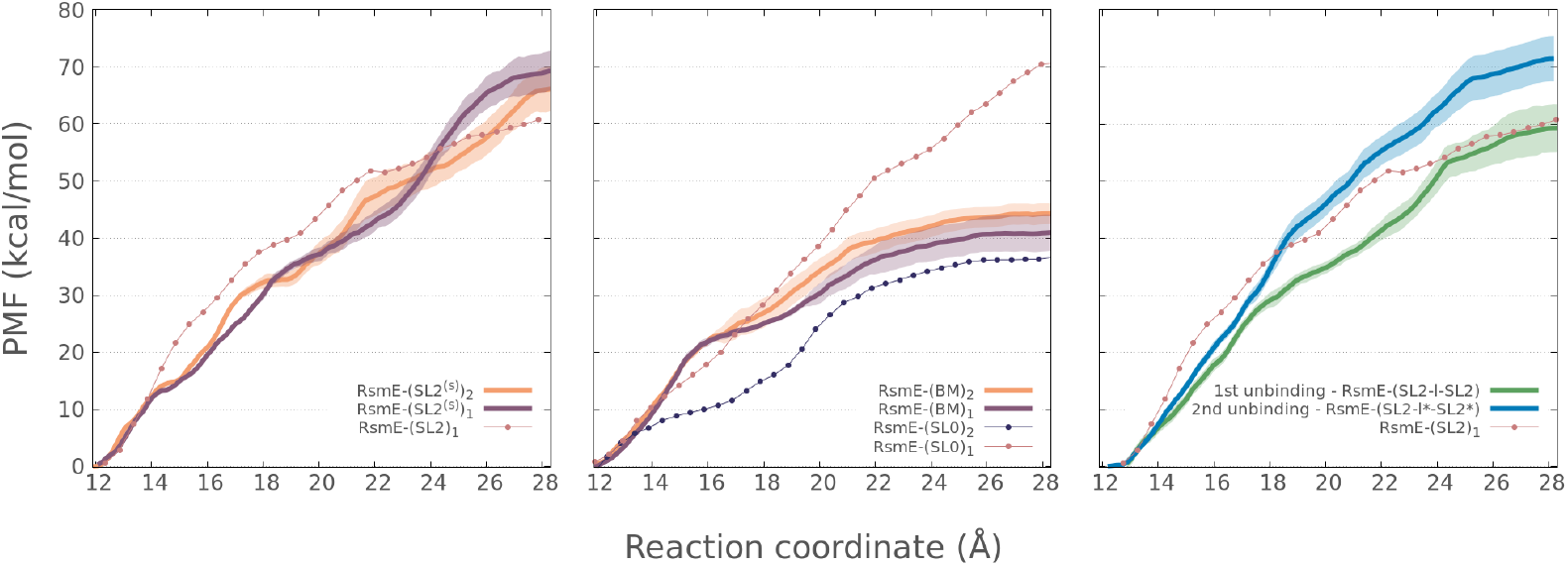
Comparison of the Potentials of Mean Force (PMFs) for the detachment of tailored RNA constructs from their complexes with RsmE. Left: SL2^(s)^ unbinding from RsmE–(SL2^(s)^)_1_ and RsmE–(SL2^(s)^)_2_; the SL2 unbinding PMF from RsmE–(SL2)_1_ is included for reference. Middle: Detachment of the 6-nt A(A)GGAC motif of SL0, corresponding to the *hcnA* fragment studied in Ref. 26, with the SL0 unbinding PMF from RsmE–(SL0)_1_ shown for comparison. Right: Detachment of each stem loop in the RsmE-(SL2^(l)^)_2_ complex; SL2 unbinding PMF from RsmE–(SL2)_1_ and RsmE–(SL2)_2_ is included for reference.

To complete the analysis, we also computed the number of contacts between RsmE and SL2^(s)^ in both, RsmE-(SL2^(s)^)_1_ and RsmE-(SL2^(s)^)_2_. The probability distributions for these contact numbers are presented in the left panel of Fig. 7, together with those obtained for the full-length SL2. It is observed that the distributions for RsmE-(SL2^(s)^)_1_ and RsmE(SL2^(s)^)_2_ are nearly identical, and they closely resemble that of RsmE-(SL2)_1_. These results indicate that the second SL2^(s)^ engages, on average, in the same interactions with RsmE as the first one. Furthermore, these interactions are comparable to those formed by a single fulllength SL2 bound to RsmE. Thus, the comparison not only corroborates that steric repulsion between the three lowest base pairs is responsible for the existence of two alternative FEBs in the complexes of the SL2 studied in Ref. 25, but also demonstrates that these base pairs do not provide stabilizing contacts with the protein, even in the complex containing a single stem loop.

**Figure 7.**
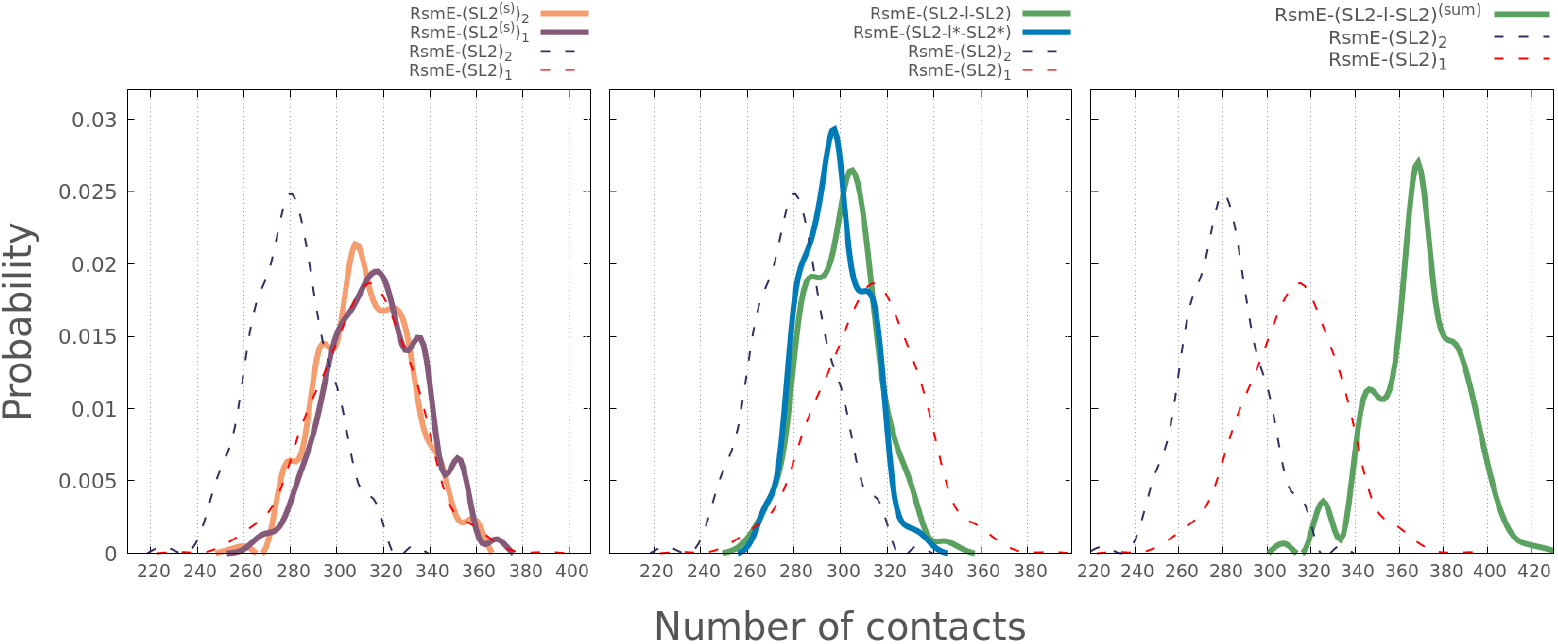
Probability distribution functions for the number of RNA-protein contacts in complexes formed by tailored RNA constructs derived from SL2. For comparison, in all cases the distributions obtained for the SL2 complexes are also shown. Left: contacts in RsmE-(SL2^(s)^)_1_ and RsmE-(SL2^(s)^)_2_. Middle: contacts in RsmE–(SL2^(l)^)_2_; since in this case the two bound stem loops are not equivalent, the distributions for both are shown separately. Right: contacts in RsmE-(SL2^(l)^)_1_. Note that, in this case, most contacts arise from the stem loop attached to the primary binding surface of RsmE. However, the other stem loop and the linker region also contribute interactions with the interface between *β*_3_ and *β*_4_ in RsmE.

### 3.4 Stem Core Boost Binding Strength

Another set of calculations with tailored RNA fragments was carried out to estimate the stem’s contribution to the binding strength. As explained in Section 2.1, we built models RsmE-(BM)_1_ and RsmE-(BM)_2_, where BM stands for A(A)GGAC, the 6-nt binding motif present in SL0, the *hcnA* fragment studied in Ref. 26. The presence of the optional nucleotide (A) in this motif has been identified as the cause of strong binding, similar to the (C) in the BM of SL2.^25^

The middle panel of Fig. 6 compares the PMFs for these two complexes with those corresponding to the whole stem loop derived from *hcnA*. It is noted that, as expected, the PMFs for both complexes, RsmE-(BM)_1_ and RsmE-(BM)_2_, are pretty similar to each other. However, in this case, the FEBs derived from them are nearly half of that corresponding to the RsmE-(SL0)_1_ complex, indicating that almost half of the interaction energy of the latter complex derives from contacts established by the stem.

### 3.5 No Stem Repulsion in Mid-Affinity Binders like SL3

Fig.4 (right panel) shows that the free energies required to remove SL3 from RsmE-(SL3)_2_ and RsmE-(SL3)_1_ are the same, within the statistical uncertainty of the calculations. This observation perfectly agrees with the experimental characterization of this stem loop that indicated there is a unique K_*d*_ for this fragment, instead of two as in the case of SL0 and SL2. In other words, for the case of SL3, the ITC experiments could be fitted with a one-site model while a two-site model was required for SL2 and the 20-nt construct derived from *hcnA*. Movie S2 illustrates the binding of SL3 to RmsE-(SL3)_1_. Contrary to what is observed in the case of SL2, we see that the attachment of the second SL3 does not alter the position of the SL3 already bound. This impression is further confirmed by the analysis of the probability distribution functions for the number of contacts, which are presented in the right panel of Fig. 5. It can be seen that the two curves are pretty similar to each other. It is also noted that the number of contacts in these two curves is significantly smaller than those in the SL2 complex, which explain the smaller FEBs and larger K_*d*_s of the SL3 complexes.

In Ref. 25, SL3 was classified as a mid-affinity binder of RsmE. This property is shared by stem loops containing the 5-nt binding motif AGGAX. In contrast, the strong binders SL2 and the *hcnA* fragment possess a 6-nt motif, A(X)GGAX. The picture that emerges from the simulation results presented so far is that, due to the weaker interaction of the SL3 binding motif, its stem remains farther from RsmE than those of SL2 and *hcnA*. The most evident consequence of this is the smaller number of RNA–protein contacts observed for SL3. A less obvious, yet important consequence, is that the lower base pairs of SL3 in RsmE-(SL3)_1_ do not occupy the space required by the lower base pairs of the incoming SL3 during the formation of RsmE-(SL3)_2_. Accordingly, the position of the already bound SL3 remains unperturbed upon binding of the second one, resulting in very similar numbers of contacts, binding free energies, and dissociation constants for the singly and doubly occupied SL3 complexes.

### 3.6 Entropic effects underlie the stronger binding of the linker-containing construct

The right panel of Fig. 6 shows the PMF profiles corresponding to the detachment of each stem loop from the RsmE–(SL2^(l)^)_2_ complex. As observed previously for the RsmE–(SLX)_1_ and RsmE–(SLX)_2_ complexes when SLX equals SL0 or SL2, the two unbinding events require different amounts of free energy. Specifically, detaching SL2_b_ when both stem loops are bound to their sites on RsmE requires approximately 60 kcal/mol, whereas removing the remaining stem loop demands an even higher energy (about 71 kcal/mol). Interestingly, the 60 kcal/mol value matches the energy needed to dissociate SL2 from the RsmE–(SL2)_1_ complex. Based on this observation and the discussions presented in previous sections, one might expect that each SL2 in the RsmE–(SL2^(l)^)_2_ complex forms a similar number of contacts with the protein as SL2 in RsmE–(SL2)_1_. However, the central panel of Fig. 7, which compares the distributions of protein–RNA contacts in RsmE–(SL2^(l)^)_2_, RsmE–(SL2)_1_, and RsmE–(SL2)_2_, shows that this straightforward expectation does not hold.

Before discussing the contact distributions, one point must be clarified. In the RsmE–(SLX)_2_ complexes, the two stem loops are strictly equivalent, whereas in the RsmE–(SL2^(l)^)_2_ complex they are not. This asymmetry arises because the linker region is not symmetric and connects the 3’ end of one stem loop to the 5’ end of the other. Due to this lack of equivalence, we plotted the two distributions separately in Fig. 6. As shown in the figure, the two distributions are pretty similar to each other and fall between those of RsmE–(SL2)_1_ and RsmE-(SL2)_2_. Based on this, one would expect the FEB of this stem loop to be lower than FEB_1_ of SL2. Instead, as mentioned in the previous paragraph, they are nearly the same. The discrepancy becomes even more evident when considering that, at the end of this first unbinding event, the outgoing stem loop remains partially in contact with the lower part of RsmE (see Fig. S5). Thus, although the original contacts of the outgoing SL2 are already lost at this stage, new contacts were established, further reducing the energetic cost of the first unbinding step. The breakdown in the previously observed correlation between the number of protein-RNA contacts and the binding free energy suggests that, in this case where the two stem loops are covalently connected via a single-stranded linker, additional contributions come into play that are not significant when the stem loops are separated.

Since the energetic (or enthalpic) contributions that correlate with the number of contacts lost during unbinding cannot account for the trends observed in the computed binding free energies, we turned our attention to the entropic contributions. Our analysis was also guided by observations from the movies illustrating the unbinding events. At the end of the unbinding from the RsmE–(SL2)_2_ complex, the detached SL2 explores a large portion of the configurational space, so that the energetic cost of the unbinding is partially compensated by an entropic gain. In contrast, during the first SL2 unbinding from RsmE–(SL2^(l)^)_2_, the linker strongly restricts the motion of the departing SL2, leading to a smaller entropic gain.

Specifically, the unbinding of all individual SLX discussed so far involves a similar (positive) entropy change. Therefore, variations in the free energy of the process,

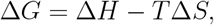

can be reasonably explained by the energetic (enthalpic) contributions, which correlate with the number of protein–RNA contacts. In contrast, for the RNA construct containing the linker segment, a smaller Δ *S* is expected, which would result in comparable Δ *G* values despite having fewer protein–RNA contacts (i.e., a smaller Δ *H*).

As detailed in Sec. 2.5, we quantified these differences using the quasi-harmonic approximation to estimate Δ *S*_SL2_ and 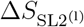. From these values, the entropic contributions to each unbinding event were determined as *T* Δ *S*_SL2_ = 349.4 kcal/mol and 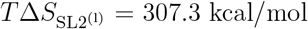. Thus, 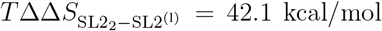, demonstrating that the entropy gain upon unbinding of the construct containing the linker is significantly smaller. This finding provides a qualitative though robust explanation for the breakdown in the previously observed correlation between the number of protein-RNA contacts and the binding free energy.

To conclude this section, we note that the removal of the remaining SL2 from the RsmE–(SL2^(l)^)_1_ complex exhibits the largest FEB among all the processes analyzed in this work. This observation can be rationalized by considering that this complex also presents the highest number of protein–RNA contacts among all the systems studied, as shown in the right panel of Fig. 5. In this case, contacts are not only formed by the SL2 bound to its canonical binding site on RsmE, but also by the detached SL2, which interacts with the lower part of the protein. Furthermore, the linker region itself establishes approximately ten contacts, on average, with the protein.

## 4 Conclusions

RNA molecules are inherently dynamic entities that populate a wide ensemble of conformations. This conformational diversity is a fundamental aspect of their function, as it underlies their ability to recognize diverse partners, regulate interactions, and adapt to changing environments. Experimental strategies often rely on dissecting large RNAs into smaller fragments to facilitate analysis. However, due to the intrinsic structural plasticity of RNA, this divideand-conquer approach must be applied with caution, as the behavior of the whole RNA molecule emerges from cooperative and context-dependent interactions that do not exist in the isolated components. The comparison between the results reported in Ref. 25 and 24 provides a great example of this.

Molecular dynamics (MD) simulations are a well-established and widely used approach for probing the molecular underpinnings of biomolecular function. Although originally developed to investigate the behavior of well-structured proteins, the scope of MD has expanded substantially over the past decades, enabling the study of more challenging and conformationally diverse systems such as RNA. In this work, we report the results of MD simulations performed on experimentally characterized RNA fragments^25^ and on in silico–designed RNA constructs tailored to identify the molecular determinants of their function. The main outcomes of these simulations are summarized below.

- The binding free energy of stem loops derived from RsmZ or *hcnA* to RsmE has significant contributions from protein–RNA interactions involving nucleotides located in the stem.
- The stem of strong binders, such as SL2 and the fragment derived from *hcnA*, locate closer to the protein surface than those of middle-strength binders such as SL3.
- In the complexes between RsmE-(SLX)_1_, with SLX equal SL2 or the SL derived from *hcnA*, the close proximity between the molecular partners is such that the three base pairs at the base of the stem occupy the space required by the second stem loop in the RsmE-(SLX)_2_ complex. This fact explains the anti-cooperative effect discussed in Ref.
- 25. We note, however, than the effect is mostly an artifact of the construct employed in the experiments (and also in our computational models), since two of the three base pairs were added to stabilize the stem.
- Simulations carried out with the model containing two SL2s connected by a linker region revealed the crucial role of this part of the molecule.
- Considering the binding of the (SL2^(l)^)_2_ to RsmE as a two stage process, the US simulations probed that the first binding is favored because the increased number of protein-RNA contacts in the intermediate complex, which not only include those between the bound SL and its canonical binding site on RmsE, but also contacts between the other SL and the linker region with non-canonical sites in the protein.
- Due to the interactions mentioned above, this intermediate state has reduced entropy (compared to the situation in which the SL2 outside its binding site is free to move). Consequently, the entropic penalty for the remaining binding event is smaller, favoring formation of the fully bound state.
- In summary: having a linker region provides an enthalpic contribution to the union of the first SL2 and an entropic contribution to the second one. This finding a simple explanation for the increased binding strengh of the stem loops in the whole RsmZ molecule than the isolated fragments.

