## Supplementary Information for "Towards the Rational Design of RsmE Small-RNA Binders: Insights from Molecular Dynamics Simulations"

### Supporting Information

#### Umbrella Sampling (US) convergence test

Alternative methodologies were employed to assess the accuracy and consistency of the Potential of Mean Force (PMF). These are listed and described below:

- One of them involves computing the PMF by means of three alternative methodologies that remove the bias from the distributions obtained in the US simulations:

the Weighted Histogram Analysis Method (WHAM),[1] the Dynamic Histogram Analysis Method (DHAM)[2] and the Bennett Acceptance Ratio (BAR).[3] For WHAM and DHAM, the reaction coordinate (RC) was binned in segments of 0.025 Å. In all cases, we used 700000 iterations for the WHAM algorithm since we had checked they were enough for convergence. For the BAR algorithm instead, we employed 500 iterations to compute the PMF difference between adjacent distributions. To implement these algorithms, we used our own FORTRAN codes that are available in our Github repository: `https://github.com/ufquilmes/RsmE/blob/main/pmf.f`.

- In a second test, we divided the whole data into three sets of equal size, and calculated the PMF (via the BAR method) for each of them. The standard deviation of these profiles were used to estimate the statistical uncertainty of the global profile.

- In a third assessment, we computed the biased probabilities for each US window in two alternative ways, and contrasted both results. The biased probabilities $P_i(\mathrm{RC})$ are those directly obtained from the US simulations in window $i$. Probabilities $\mu_i(\mathrm{RC})$ are estimated from the unbiased distribution obtained with WHAM by adding the effect of the bias potential. The consistency between the two distributions was evaluated with the symmetric Kullback-Leibler divergence test,

$$S_i = \frac{1}{2}\ D(P_i, \mu_i) + \frac{1}{2}\ D(\mu_i, P_i)\,, \tag{1}$$

where,

$$D(f, g) = \sum_{k=1}^{N} f(RC_k)\ ln\left(\frac{f(RC_k)}{g(RC_k)}\right). \tag{2}$$

In Eq. 2 $N$ represents the number of bins employed in a discretized representation of the probability densities $f(\mathrm{RC})$ and $g(\mathrm{RC})$, while $\mathrm{RC}_k$ is the value of the random variable at the center of bin $k$. The lower the value of $S_i$, the better agreement between the two distributions. We observed that all values considering the whole group of studied models, were below 0.005 indicating consistency between the raw data from the US-simulations and the results afforded by means of the BAR methodology.

### 1 List of Figures

- Figure S1: Tests afforded to evaluate the consistency of US simulations for the binding event between RsmE and all the *hcnA* fragments studied in this work. PMFs obtained with WHAM, DHAM and BAR are compared. Equivalent results take place for all the remaining models here studied.

- Figure S2: Tests afforded to evaluate the consistency of US simulations for the binding event between RsmE and all the *hcnA* fragments studied in this work. Comparison of the independent PMF calculations obtained considering the sampled data divided into three consecutive thirds. Equivalent results take place for all the remaining models here studied.

- Figure S3: Symmetric Kullback-Leibler divergence test to check US simulation results for the binding event RsmE and all the *hcnA* fragments studied in this work. Equation 1 presents the formula for computing this parameter as a function of the reaction coordinate $\chi$. Equivalent results take place for all the remaining models here studied.

- Figure S4: QHA Entropy estimations with respect to the number of individual trajectories included in the concatenated one.

- Figure S5: Pictoric representation for the RsmE-(SL2-l*-SL2*) complex.

### 2 List of Movies

- Movie S1. Binding process of SL2 to the RsmE-$(SL2)_1$ complex.

- Movie S2. Binding event between

### 3 Fortran files

- “pmf.f”: Fortran code implemented to estimate the Potential of Mean Force of the unbinding event by means of WHAM, DHAM and BAR, as well as to asses their consistency and statistical uncertainty.

### 4 Figures

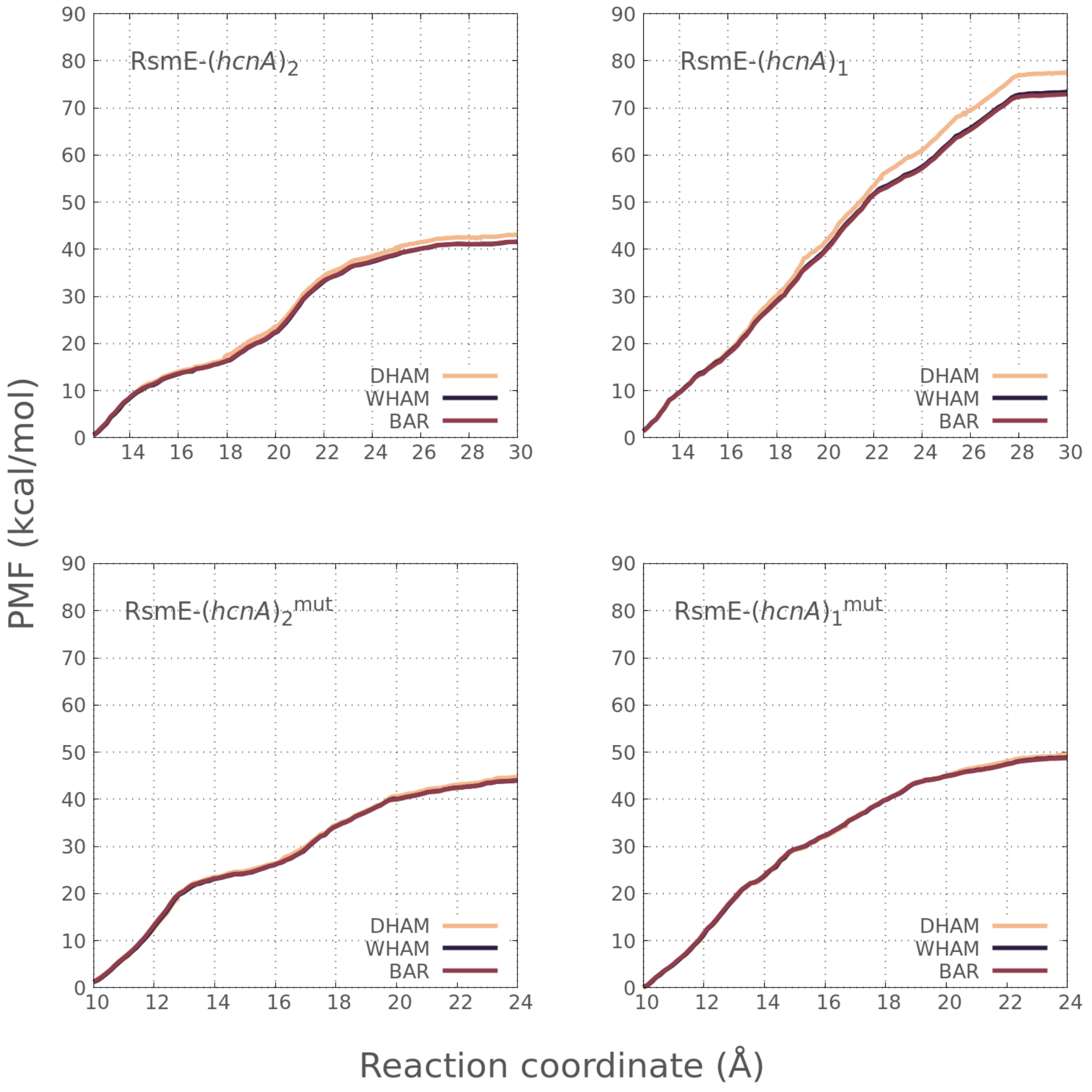


Figure S1: Comparison of the PMFs calculated with WHAM (blue), DHAM (orange) and BAR (red) for the four models studied in this work by means of Umbrella Sampling calculations.

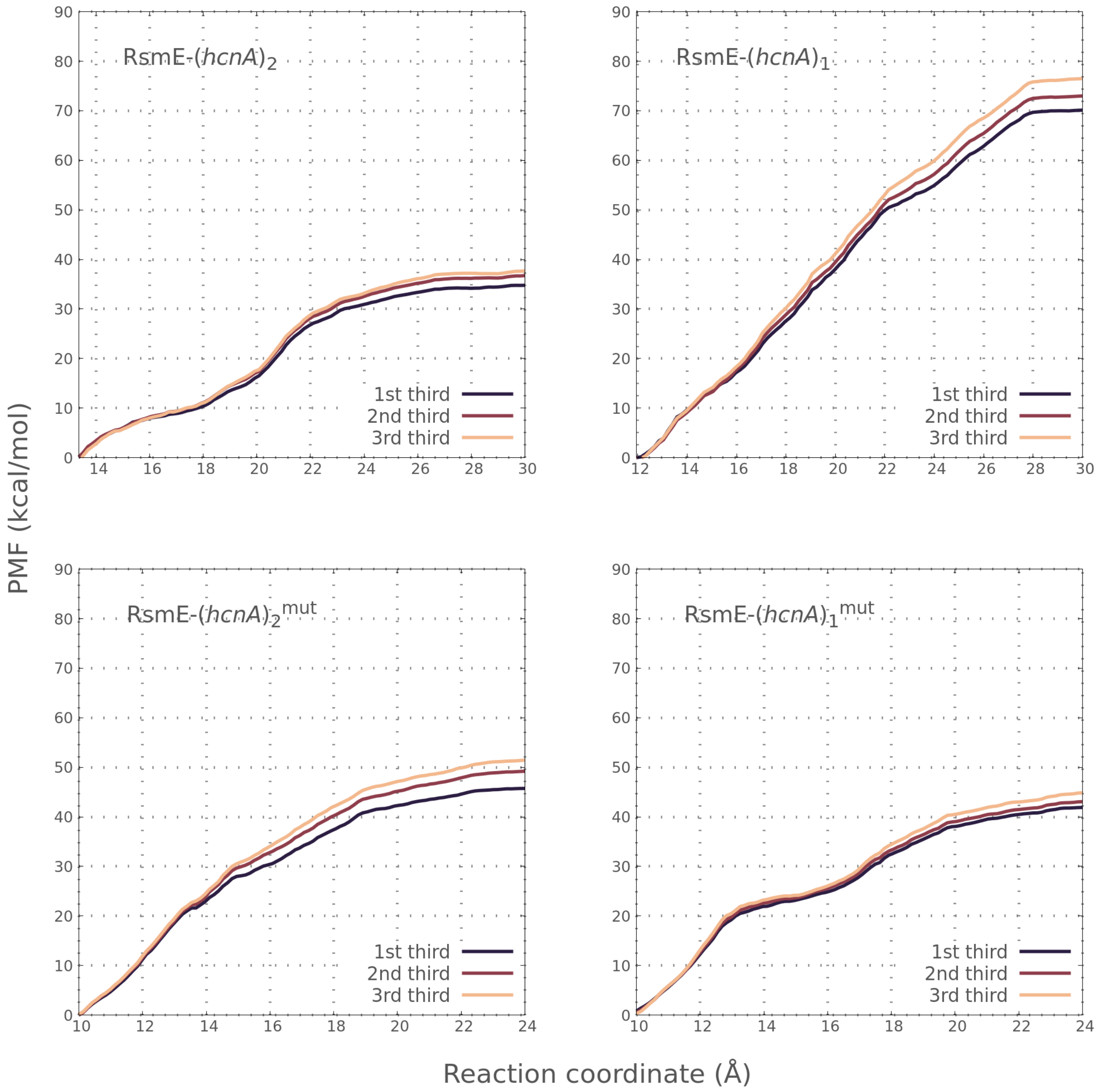


Figure S2: Comparison of the PMFs calculated with three alternative data sets obtained by dividing the whole set in three equal parts. Two insets zooming in segments at different parts of the profile are presented to highlight the similarity between the three PMFs.

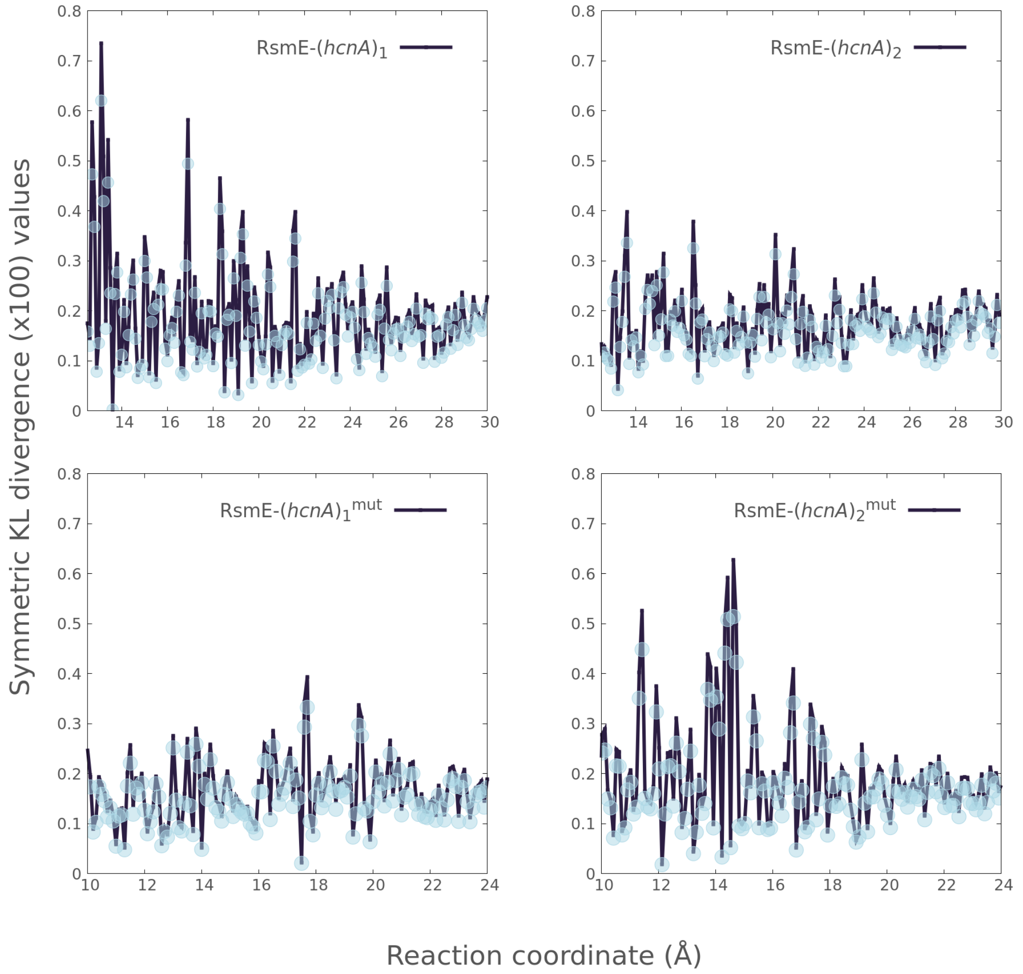


Figure S3: Panel A illustrates the sKL-divergence (Eq. 1) as a function of the RC. Panel B compares the PMF computed with alternative data sets obtained by dividing the whole set in three thirds while panel C does it when the WHAM (red), DHAM (blue) and BAR (green) methodologies are implemented considering the whole data.

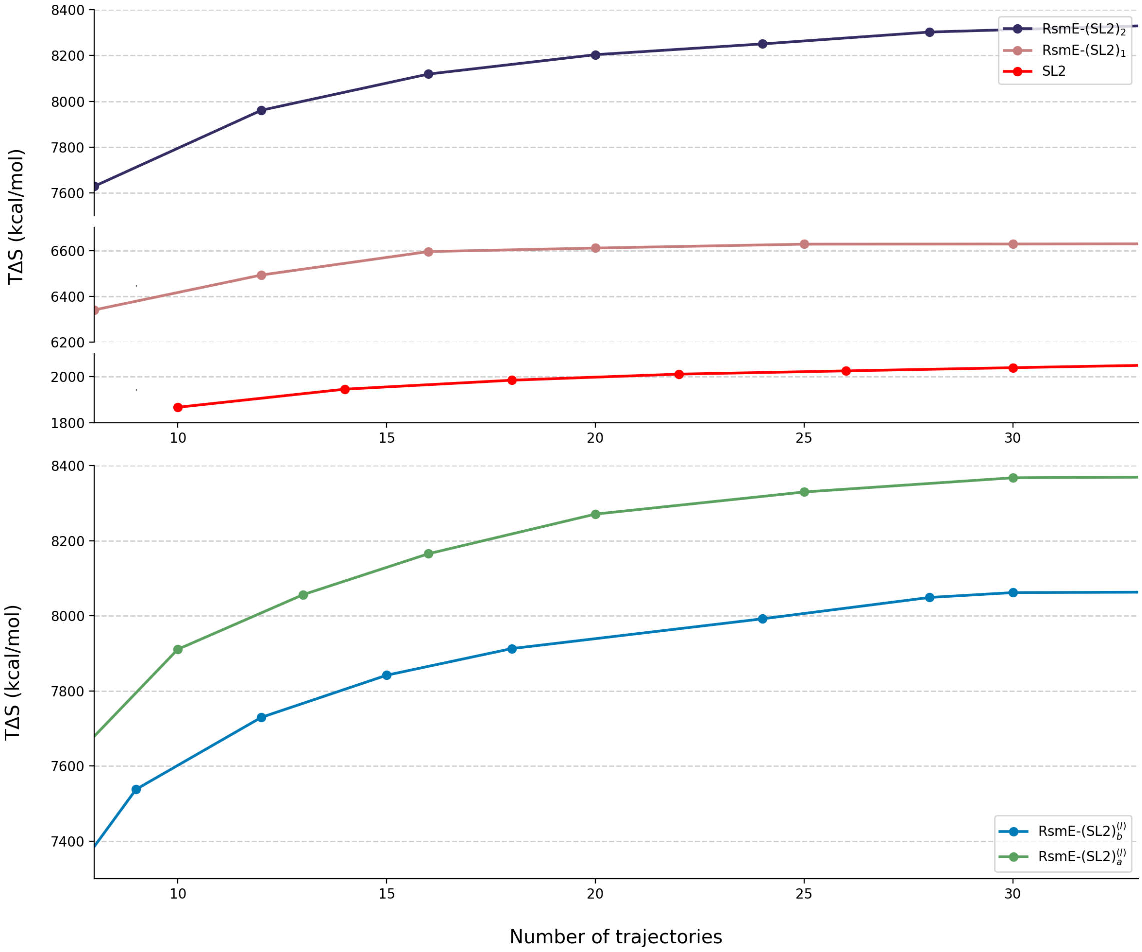


Figure S4: Convergence of the entropy estimations with respect to the number of individual trajectories included in the concatenated one (See Eq. 2).

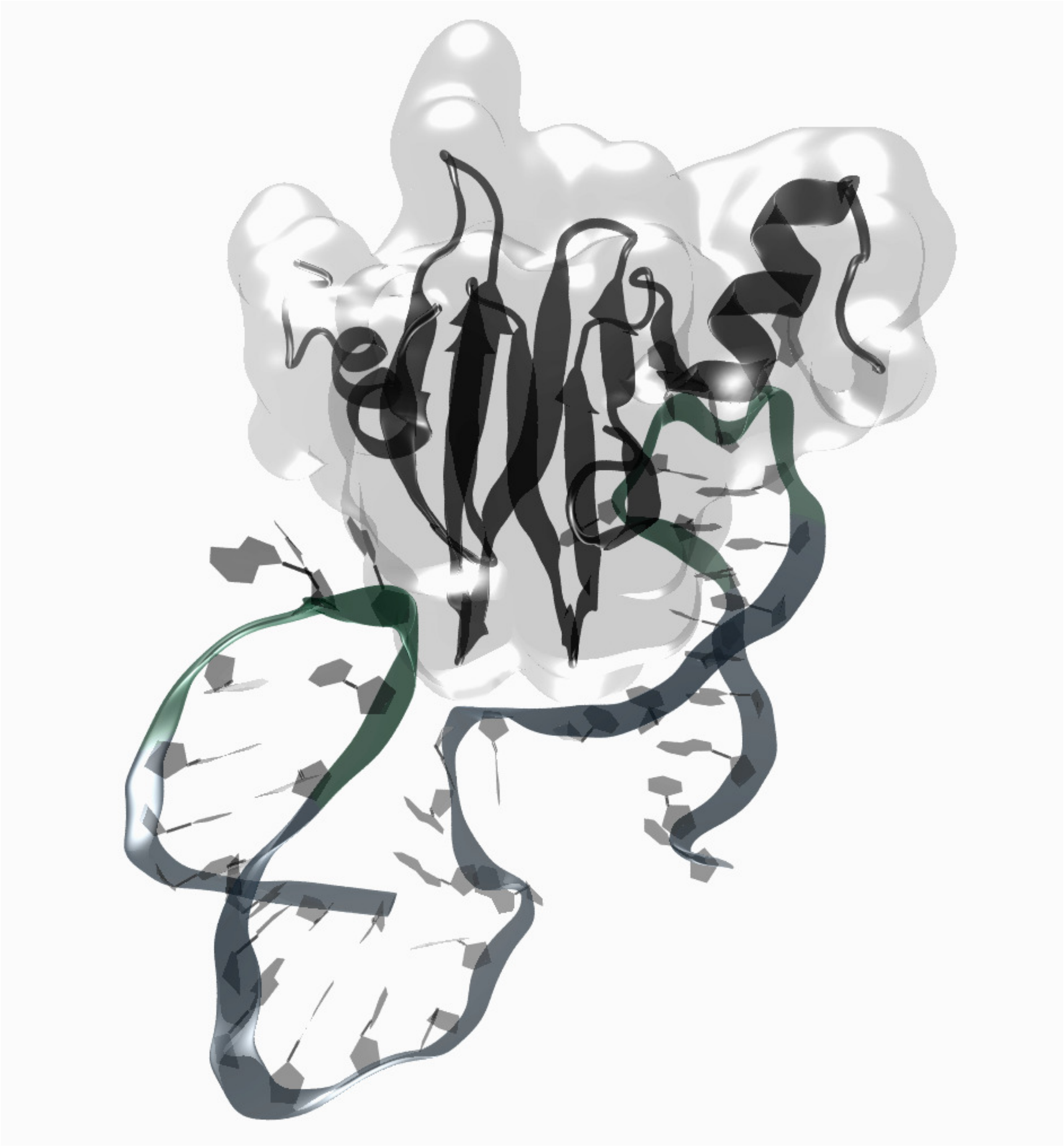

Figure S5: Pictoric representation of the RsmE-(SL2-l*-SL2*) complex.
